# Effects of Standing on Spinal Alignment and Lumbar Intervertebral Discs in Young, Healthy Individuals Determined by Positional Magnetic Resonance Imaging

**DOI:** 10.1101/522565

**Authors:** Christian I. Weber, Ching-Ting Hwang, Linda R. Van Dillen, Simon Y. Tang

## Abstract

Traditional diagnostic imaging of the spine is performed in supine, a relatively unloaded position. However, the spine is subjected to complex loading environments in daily activities such as standing. Therefore, we seek to quantify the changes from supine to standing in the spines of young, healthy individuals in standing using a positional MRI system. This is an observational study that examined the changes in the spine and individual intervertebral discs (IVDs) during supine and standing of forty healthy participants (19 males / 21 females) without a history of low back pain. The regional lumbar spinal alignment was measured by the sagittal Cobb angle. Segmental IVD measurements included the segmental Cobb angle, anterior to posterior height (A/P) ratio, and IVD width measured at each L1/L2 - L5/S1 levels. The intra-observer intra-class correlation (ICC) consistency model showed values for measurements ranged from 0.76-0.98. The inter-observer ICC values ranged from 0.68-0.99. The Cobb angle decreased in standing. The L5/S1 segmental Cobb angle decreased in standing. The L2/L3 and L3/L4 A/P ratios increased and the L5/S1 A/P ratio decreased in standing. No differences in IVD width were observed from supine to standing. This study examined the regional lumbar spinal alignment and segmental IVD changes from supine to standing in young, healthy individuals without LBP using pMRI. In developing and validating these measurements, we have also established the normative data for healthy, asymptomatic population that could be useful for other investigations examining how individuals with spinal or IVD pathologies may adapt between supine and standing.

## Introduction

The degeneration of the intervertebral disc (IVD) is a significant contributor to low back pain (LBP) (39). Despite the association between IVD degeneration and LBP, the correspondence between the clinical presentation of LBP and IVD imaging findings is quite poor (35). One possibility for this lack of specificity may be that most diagnostic imaging of the spine is performed in supine, a minimally loaded position (5, 45). An individual in standing generates lumbar intradiscal pressures approximately five times greater than in supine (31) and complex multiaxial forces (32) that result in different tensile and shear deformations across the IVD (3, 43). Compressive loading similar to that experienced during standing is critical for maintaining spinal curvature (44). Thus, examining the spine in standing may be more functionally and clinically relevant than in supine.

Magnetic Resonance Imaging (MRI) is a commonly used radiographic technique for the imaging of the spine and for the visualization of hydrated tissue structures such as the IVDs. Positional MRI (pMRI) with an open magnet configuration enables the imaging of the human participants in different positions (4, 18, 19); thus, the pMRI provides the opportunity to observe the spinal structures in loaded positions other than supine (6, 18, 36, 37). Examining the lumbar spine in an upright position using a pMRI has improved the specificity in discriminating acute and chronic LBP populations (42) and increased the reliability of observing stenosis and IVD degeneration (12). Prior studies have utilized pMRI to examine spinal adaptations between sitting to standing (20), supine to standing in athletes (29) and patients with lumbar spinal stenosis (27), and changes of the dural sac in the lumbar spine due to posture (16). But none of these studies explored differences of segmental structural measurements from supine to standing in a young, healthy population without a history of LBP (‘back-healthy’). Back-healthy individuals have been shown to be susceptible to LBP symptoms in prolonged standing with a high rate of developing chronic LBP (9, 10, 28, 33, 34, 40). Thus examining the standing-induced adaptations in the back-healthy population could provide predictive information prior to development of spinal pathologies. The pMRI can leveraged to observe adaptations across positions of the lumbar spine using regional alignment measurements (sagittal Cobb angle) and segmental lumbar measurements such as segmental Cobb angle, anterior/posterior height (A/P) ratio, and IVD width. Therefore, the objectives of this study are to 1) investigate the effects of standing on the spinal alignment and individual lumbar IVDs of young, back-healthy human participants ranging from 18 to 30 years of age using pMRI, and 2) determine whether there are sex-specific differences in these measurements due to the variations in spinal alignment between males and females (15, 47). We hypothesized that 1) the sagittal Cobb angle would be significantly different between supine and standing, 2) individual IVDs across the lumbar spine would adapt differentially to standing, and 3) the change in segmental Cobb angle, A/P ratio, and IVD width at each lumbar level from supine to standing would be different between males and females.

## Material and methods

### Participants

The study included forty participants (19 male/21 female) between 18-30 years of age and body-mass index (BMI) < 30kg/m^2^ (Table 1). Participants were recruited through posted flyers and distributed emails to the community and local universities in the St. Louis metropolitan area. Participants were excluded during screening if they reported any history of LBP. LBP was defined as pain in the lumbar region greater than 2 on a 0-10 verbal numeric rating scale that lasted at least 24 hours that resulted in one or more of the following: (1) some type of medical intervention (e.g., physician, physical therapist, chiropractor); (2) three or more consecutive days of missed work or school; (3) three of more consecutive days of altered activities of daily living. Exclusion criteria also included a prior diagnosis of diabetes, anxiety, depression, employment in a job that involved standing for greater than 4 hours per day or standing in one place for more than 1 hour per day during the last 12 months, consumption of caffeinated drinks > 25 per week, consumption of alcoholic drinks > 10 per week, or smoking cigarettes > 15 per day. All participants read and signed an informed consent form in accordance with the Human Research Protection Office at Washington University School of Medicine. The participants were instructed to avoid non-habitual strenuous physical activity (e.g., running, weight lifting) for 24 hours prior to imaging with no rescheduling required. All imaging was performed in the afternoon after 12PM to minimize diurnal variations (26).

**Table 1:** Characteristics of the participants presented as the mean ± standard deviation. Age (*p* = 0.81) and BMI (*p* = 0.35) are not statistically significant between sexes.

|  | **Male (19)** | **Female (21)** |
| --- | --- | --- |
| Age (y) | 24.7±3.5 | 24.9±2.1 |
| Height (cm) | 179.2±7.4 | 164.6±7.8 |
| Weight (kg) | 73.3±9.0 | 60.4±5.2 |
| BMI (kg/m<sup>2</sup>) | 23.0±2.4 | 22.3±1.9 |

### Data collection

Images of the lumbar spine (L1-S1) were obtained using the 0.6T Open UPRIGHT® MRI (Fonar, New York, NY) system. A 3-plane localizer was used to acquire sagittal T2 weighted images (repetition time = 610 ms, echo time = 17 ms, field of view = 24 cm, acquisition matrix = 210 × 210, slice thickness = 3 mm, no gap, scan duration = 2 min) (37). This sequence was optimized for reducing scan time and motion artifacts [12–13]. A wood plank was placed adjacent to the quad-planar coil on the MRI table to provide a continuous flat surface for the lumbar spine (Fig. 1). The participant entered the scanner facing forward with his/her back against the table. A pillow was placed behind the head to provide support to the neck.

**Figure 1:**
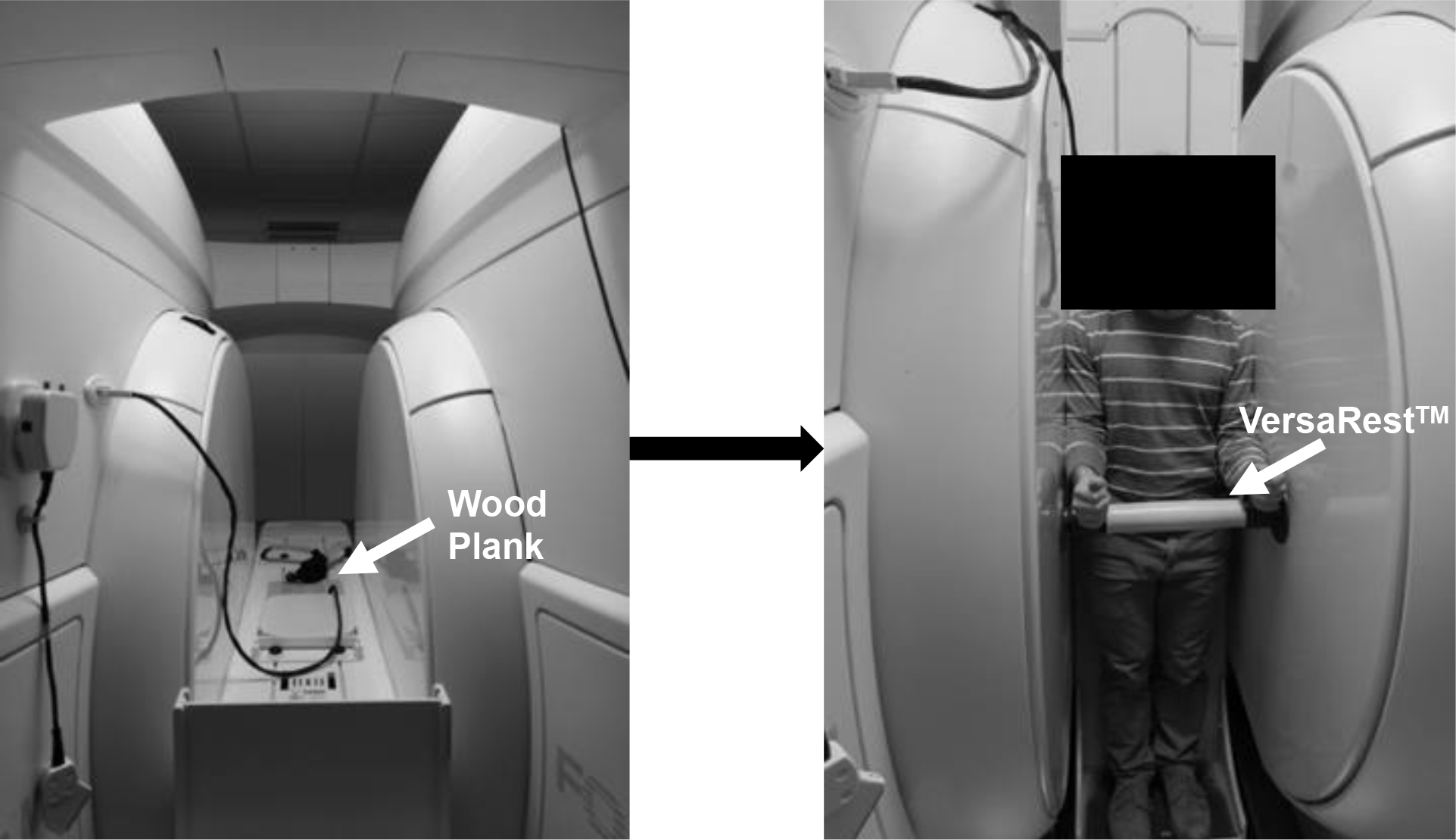
The pMRI with the table in supine (left) and standing (right) configurations. A wood plank placed behind the participant to provide a continuous flat surface for the lumbar spine and is indicated by the white arrow in the left panel. A VersaRest™ device to support the arms is indicated by the white arrow in the right panel of the figure.

The table was adjusted to a horizontal position of 180 degrees. The participant was positioned in supine for 10 minutes prior to the first scan. The MRI table then was moved to a vertical position with a 84 degree table tilt where participants were then standing. The table tilt of 84 degrees helped to stabilize the participant and prevent motion artifacts during the imaging in standing (Fig. 1). The pillow was removed, and a VersaRest™ device, an arm support, was placed in the scanner underneath the wrists 5 cm below the lateral epicondyle of the elbow (Fig. 1). Participants were told to stand normally without leaning on the sides of the magnet, back of the scanner or on the VersaRest™ during the scan in standing. A board-certified MRI technician conducted all imaging. After imaging, all images were exported as DICOM files to be analyzed on Miele (OsiriX)-LXIV (open source) (38).

### Regional Lumbar Spine Alignment

The four-line Cobb method, denoted here as the Cobb angle, was used to quantify regional lumbar spine alignment (14). The Cobb angle was measured between the inferior T12 and superior S1 endplates (Fig. 2A). This measurement was obtained in supine and standing for each participant using the mid-sagittal slice of the MR image.

**Figure 2:**
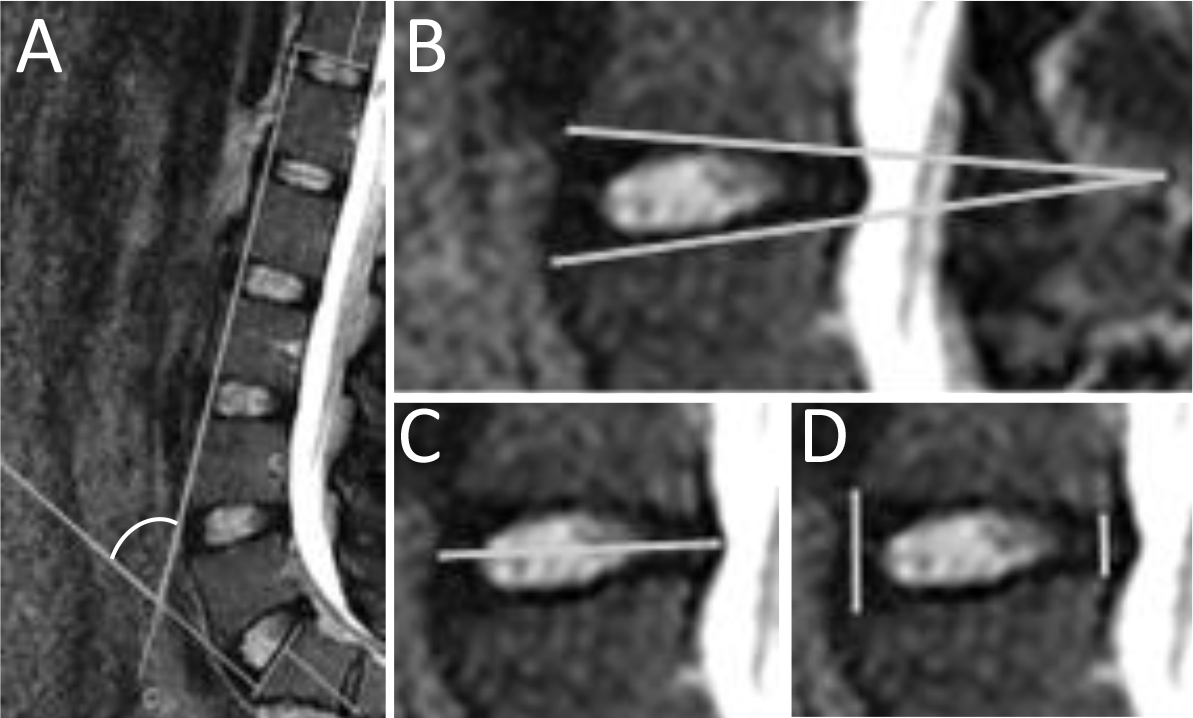
A representative image of the measurements in this study: A) The four-line Cobb angle, with the white arc indicating the angle of measurement, quantifies the regional spinal alignment. Larger Cobb angles reflect more lumbar lordosis. B) The segmental Cobb angle describes the “wedging” of the IVD. C) The anterior to posterior height (A/P) ratio quantifies the relative change of shape of the IVD. D) The IVD width quantifies the radial bulging that occurs in the IVD. All images are aligned in the same orientation with the left being the anterior side.

### Segmental IVD Measurements

The measures for each of the five lumbar IVDs, L1/L2 through L5/S1, were obtained in supine and standing for each participant using the mid-sagittal slice of the MR image. The measures included the segmental Cobb angle, the anterior to posterior height (A/P) ratio, and the IVD width (Fig. 2B, 2C and 2D). The segmental Cobb angle was calculated as the angle created by the line segment containing the anterior and posterior edges of the superior endplate with the line segment containing anterior and posterior side of the inferior endplate (Fig. 2B). The heights were calculated as the distances on the superior endplate to inferior endplate distance at the anterior and posterior sides for each IVD level. The A/P ratio was calculated by dividing the anterior height by the posterior height (Fig. 2C). The IVD width was defined as the maximum distance between the anterior and posterior bulge points of an IVD in a sagittal view (Fig. 2D).

### Reproducibility, error and uncertainty

The intra-observer (C.W./researcher) and inter-observer (researchers) reproducibility were indexed with the Intraclass Correlation Coefficient (ICC) using data from participants that exhibited the greatest range of values for regional and all lumbar segments (30). A written rubric based on anatomical landmarks (segmental Cobb angles, IVD widths, the anterior-, and posterior-heights) was used for all segmental measurements. The intra-observer analysis was performed using measured variables obtained on three distinct nonconsecutive days. Additional observers performed the inter-observer analysis based on the written rubric for regional and segmental measurements with three sets of measurements on non-consecutive days.

We quantified the error associated with image resolution and its impact on the uncertainty of IVD measurements. The resolution-based error of the MRI images was determined by moving the coordinates of each anatomical landmark in every possible direction and calculating the effect on the subsequent IVD measurements. The maximum uncertainty from resolution on the measurements using this combinatorial approach were 5.2%, 5.4%, and 1.3% for segmental Cobb angle, A/P ratio, and IVD width, respectively.

### Statistical Analyses

A three-way, repeated measures analysis of variance (ANOVA) was used to test for the factors of position, sex, and lumbar level for each of the three segmental measurements (segmental Cobb angles, A/P ratios, and IVD widths). Cobb angle was analyzed using a two-way repeated measures ANOVA to test for the factors of position and sex. Interaction terms are denoted by an asterisk (*) in Table 2 and in the results. Post-hoc comparisons were done using the Fisher’s least significant difference test. Differences are significant when the associated p-value is ≤ 0.05. Post-hoc power analysis was performed on significant results using G*Power (8).

**Table 2:** P-values for the three-way repeated measures ANOVA examining the effects of sex, level, and position and their interactions on the segmental Cobb angle, A/P ratio, and IVD width measurements.

| Factors | *p*-value |  |  |
| --- | --- | --- | --- |
|  | Segmental Cobb angle | A/P ratio | IVD width |
| Sex | 0.027 | < 0.001 | < 0.001 |
| Level | 0.004 | < 0.001 | < 0.001 |
| Position | --- | < 0.001 | < 0.001 |
| Sex*Level | --- | --- | --- |
| Sex*Position | --- | --- | --- |
| Level*Position | < 0.001 | < 0.001 | < 0.001 |
| Sex*Level*Position | --- | --- | --- |

## Results

All enrolled participants (n = 40) completed the supine and standing imaging sessions and were included in all analyses.

### Reliability of Measurements

Figure 3 shows intra-observer ICC values for each measurement. The ICC values (with the 95% confidence interval in parentheses) for the segmental Cobb angle, IVD width, and the anterior and posterior height measurements were 0.90 (0.79-0.96), 0.98 (0.95-0.99), 0.89 (0.78-0.95), and 0.76 (0.58-0.89) respectively. The inter-observer ICC agreement for segmental Cobb angle, IVD width, and the anterior and posterior height measurements were 0.89 (0.79 – 0.95), 0.99 (0.98 – 1), 0.84 (0.7 – 0.93), and 0.68 (0.46 – 0.85), respectively. The intra-observer ICC for Cobb angle was 0.97 (0.90 – 0.99), and the inter-observer ICC was 0.87 (0.61-0.97).

**Figure 3:**
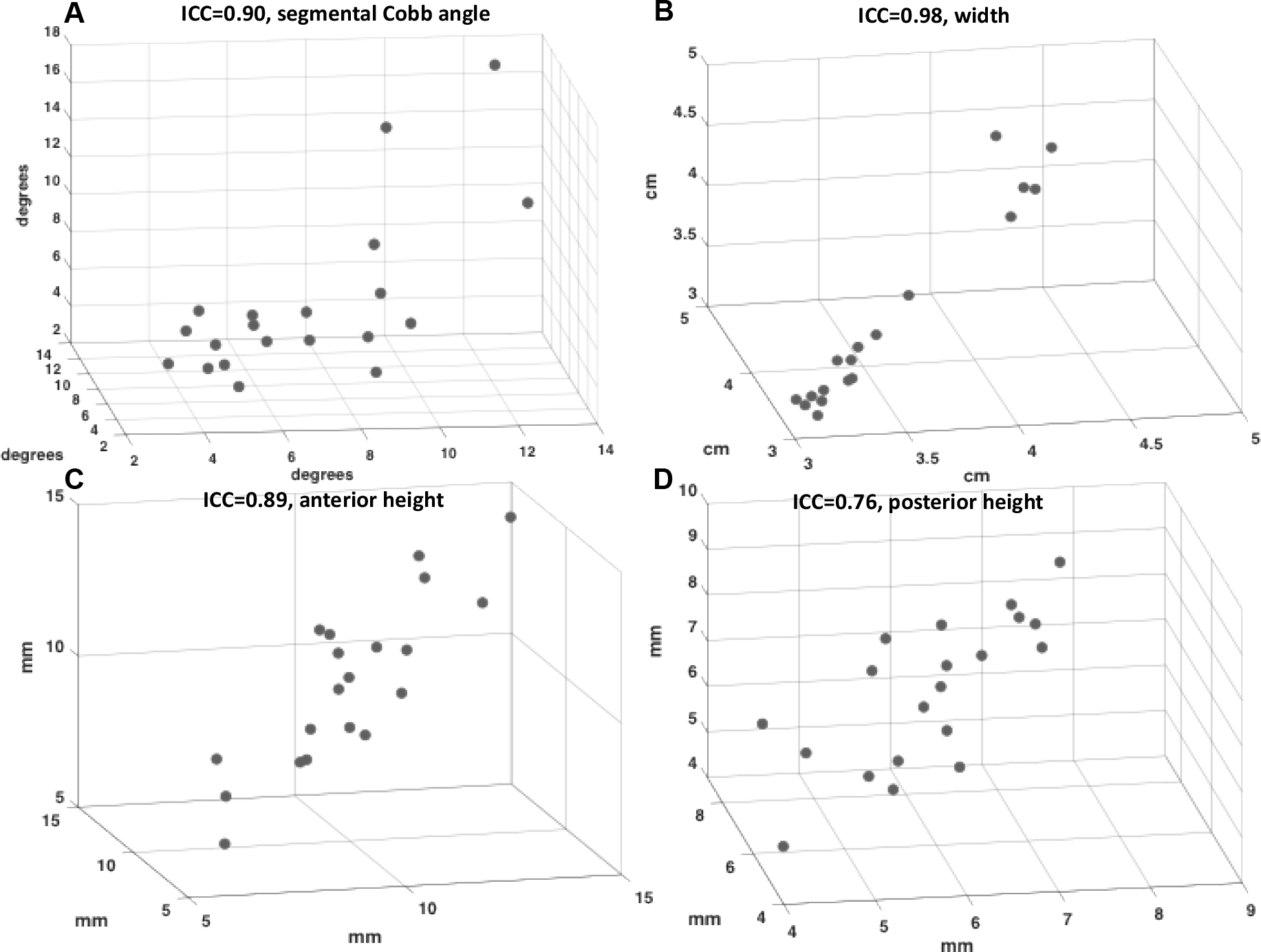
The intra-observer ICC values are shown for each measurement for individual IVD levels for A) segmental Cobb angle, B) IVD width, C) anterior height, and D) posterior height. Each axis represents an independent observer trial for the measurement. A written rubric for measurement based on anatomical landmarks (segmental Cobb angle, IVD width, the anterior-, and posterior-heights) was used. The intra-observer analysis was performed using measured variables obtained on three different nonconsecutive days.

### Regional Lumbar Spine Alignment

Position was a significant factor. The Cobb angle decreased in standing compared to supine for all participants (Fig. 4; p < 0.05).

**Figure 4:**
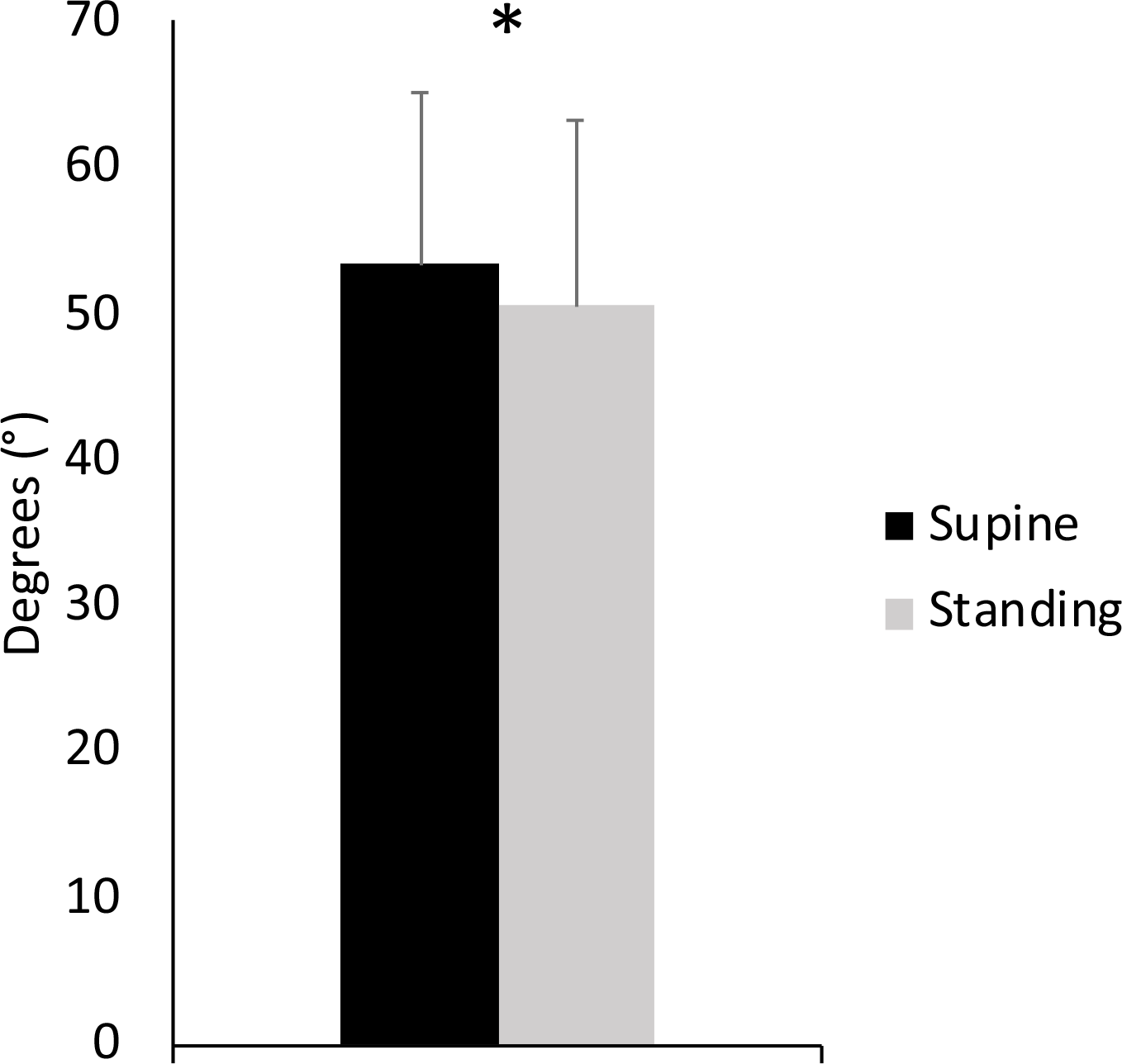
The Cobb angle for all participants in each position show that lordosis decreases in standing (* :two-way repeated measures ANOVA, *p* < 0.05). Error bars indicate standard deviations.

### Segmental IVD Measurements

#### Segmental Cobb Angle

Sex, level, and level*position all were significant factors (Table 2). Post hoc analyses revealed that compared to supine the L5/S1 segmental Cobb angle decreased in standing (Fig. 5; p < 0.05).

**Figure 5:**
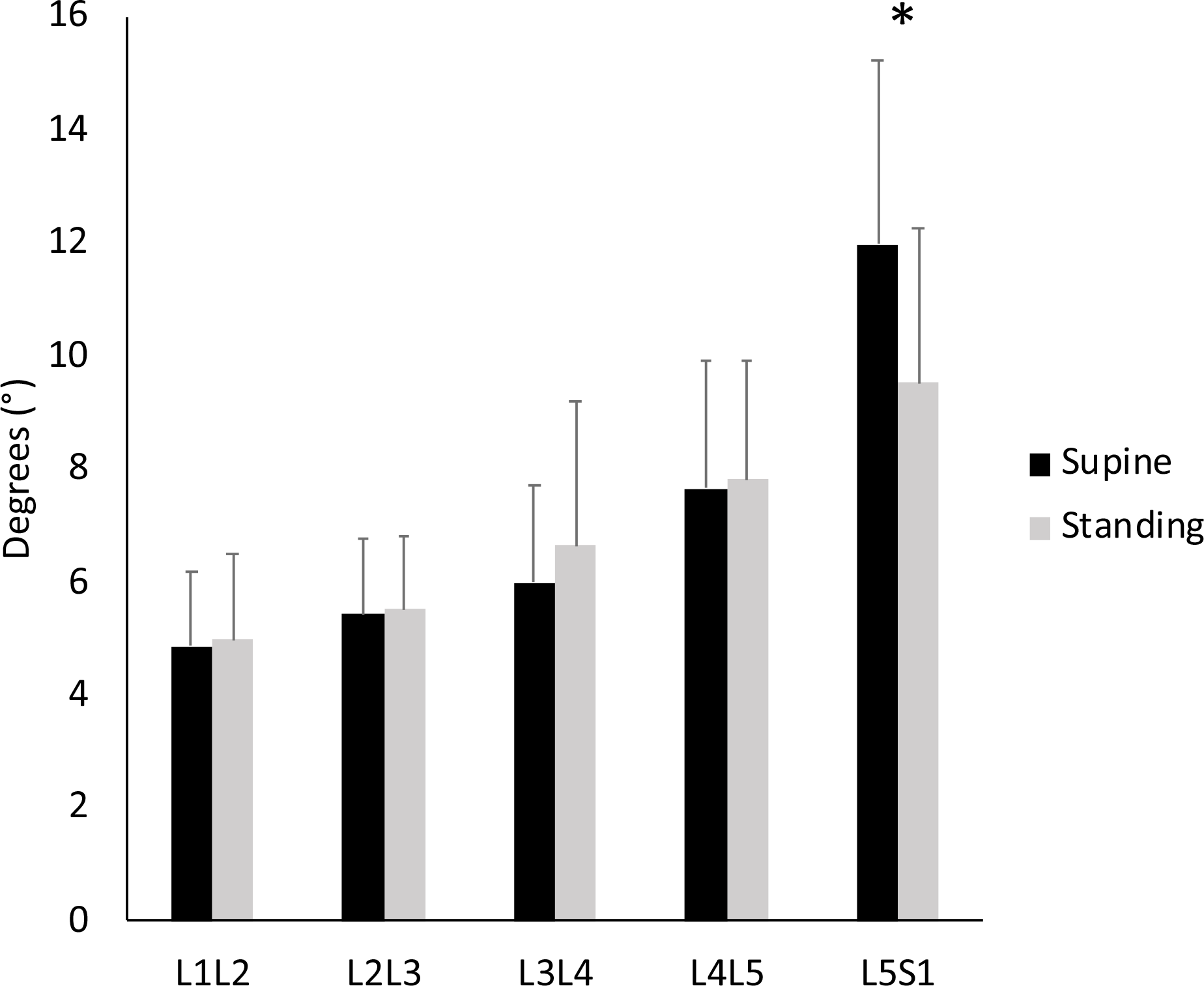
The mean segmental Cobb angles at each lumbar level for all participants in supine and in standing. The segmental Cobb angle decreased at L5/S1 in standing (* :three-way repeated measures ANOVA, *p*<0.05). Error bars indicate standard deviations.

#### A/P ratio

Sex, level, position, and level*position were significant factors (Table 2). Compared to supine the L2/L3 and L3/L4 A/P ratio increased in standing (p < 0.05), and the L5/S1 A/P ratio decreased in standing (Fig. 6; p < 0.05).

**Figure 6:**
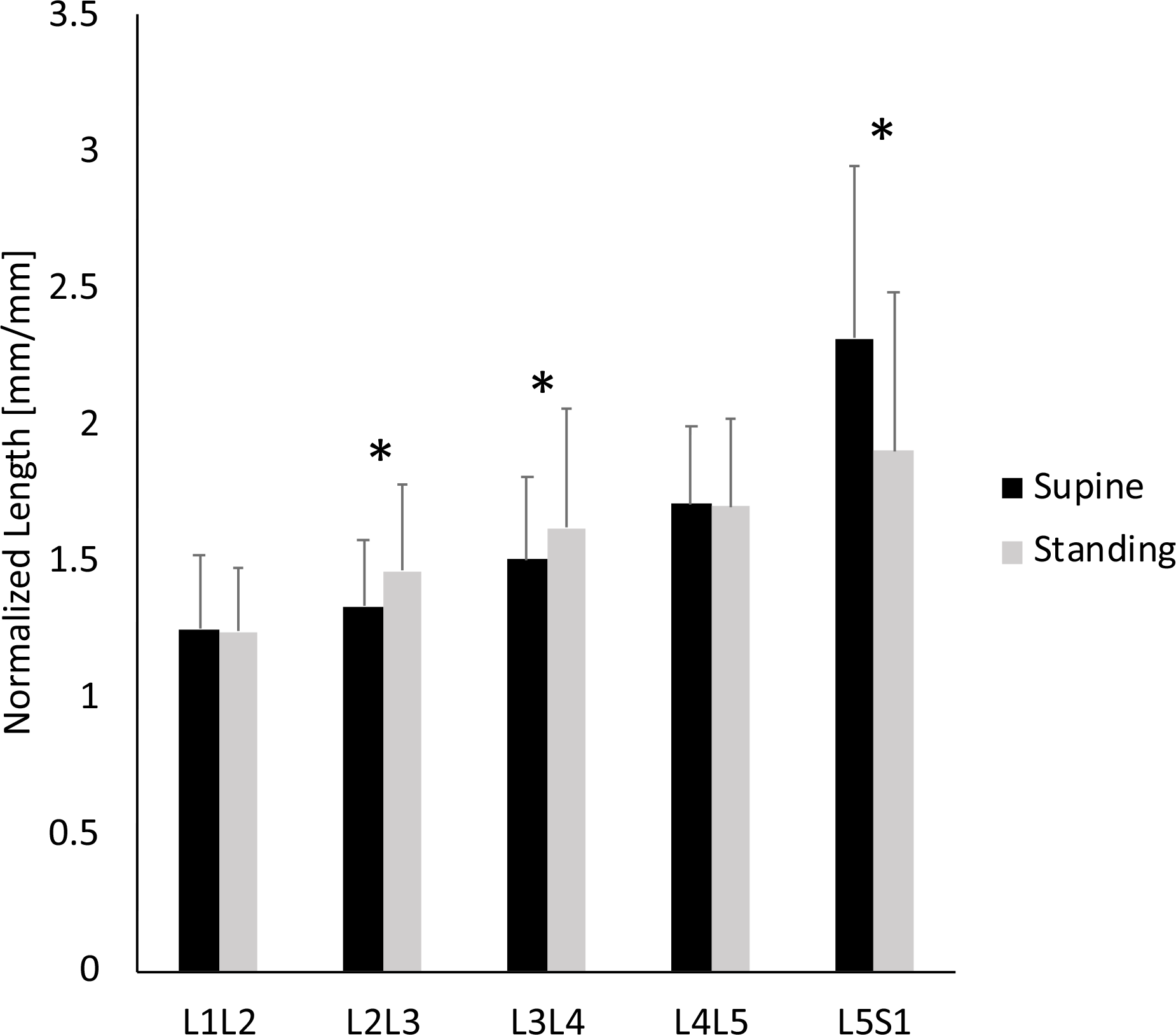
The A/P ratios show that L2/L3 and L3/L4 increases in standing, and decreased in the L5/S1. (* : three-way repeated measures ANOVA, p<0.05). Error bars indicate standard deviations.

#### IVD Width

Sex, level, position, and level*position were significant factors (Table 2). There were no significant differences in IVD width due to positions at lumbar levels L1/L2 to L4/L5 for all participants (Fig. 7). Although the L5/S1 IVD width in standing was statistically greater than in supine (p < 0.05), the margin of the difference (0.48%) was below the detection threshold per our error analyses (1.3%), and thus we did not consider this difference to be meaningful.

**Figure 7:**
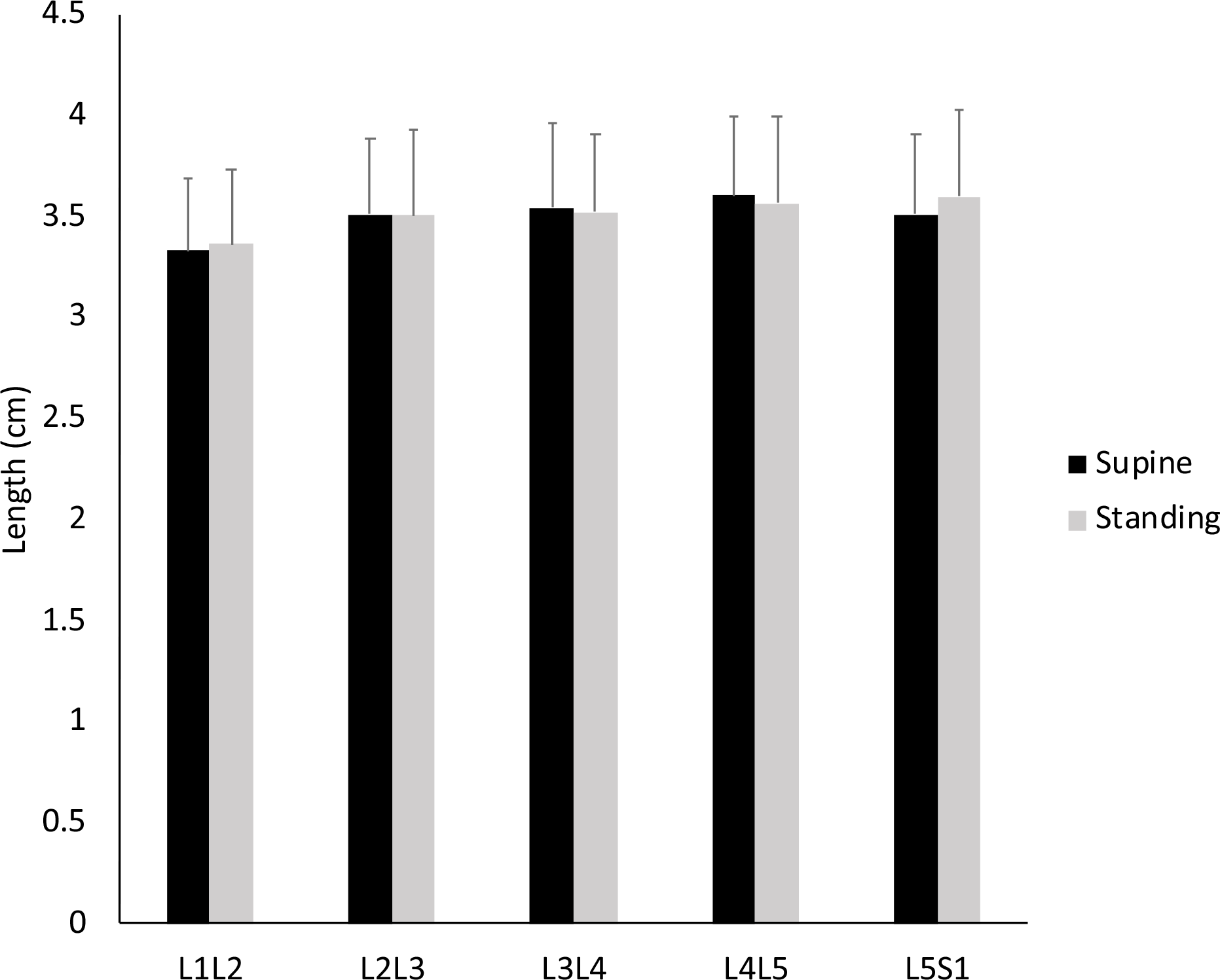
The IVD width shows no differences at all levels between supine and standing positions across participants. Error bars indicate positive standard deviations.

## Discussion

To our knowledge this is the first study to examine the segmental Cobb angle, A/P ratio and IVD width from supine to standing. In contrast to other studies that examined angular changes that included the vertebrae and the adjacent IVD (23, 46), we measured the changes between the inferior and superior vertebral endplates of individual discs (1, 17) enabling observation of IVD-specific adaptations at every lumbar level. pMRI studies on Marine populations have shown that individual IVDs adapt to external loads placed on the spine in standing (6, 37). The changes in segmental IVD measurements from supine to standing in our back-healthy group show that adaptations due to increased loading occur at multiple levels of the lumbar spine, similar to the externally loaded spine in the supine position (21). The ability of these lumbar levels to adapt may be important for the spine to distribute loads, especially when the diminished ability of individual lumbar levels to adapt to loading are concomitant with disease and degeneration (6, 22). Given that the pMRI in standing improves the detection of IVD bulging (49), the lack of bulging in these participants in standing confirms that these IVDs are indeed relatively healthy (2, 41).

The reduced lordosis of our participants in standing is consistent with the observations of another pMRI study (27). Other studies using a pMRI have shown that lordosis is not different between supine and standing with a cushion underneath their knees (16). These results are in contrast with studies done using plain film radiography which observed more lordosis in standing (7, 23, 46). The discrepancy is likely due to a pelvic tilt that has been shown to influence lumbar lordosis in standing (24), since the pelvis is strongly correlated to lumbar lordosis in supine (7). In this study, we did not directly control for pelvic tilt by placing a pillow underneath the legs in supine. Despite this, our measured Cobb angles in standing are consistent with a prior report that also included young, asymptomatic individuals (48). Further, it has been shown that externally applied loads during standing can alter the lumbar spine to be in less lordosis in pMRI (6, 36, 37). Our study did not find lordosis to be different between sexes as previously reported (15, 47). Additionally, we observed that males and females comparably change from supine to standing. Wood et al. included males and females in their study but did not specifically report on sex differences in position (46). Although we observed sex as a significant factor in the segmental measurements, there were no interactions between sex and position, suggesting though both sexes adapt to standing, they do not adapt differently. In this study, the decline of the Cobb angle of the spine in standing in a young, back-healthy group with no observed translational lumbosacral anatomies or spondylolysis/spondylolisthesis is coupled with measurable changes in the IVD across the different lumbar levels (Fig. 5, 6).

Characterizing the IVD using measurements that are based on independent landmarks is important for capturing the nuanced changes in the structure of the IVD (12, 16, 42). Using the pMRI system, we demonstrate that it is possible to determine and reliably characterize the regional lumbar alignment and individual lumbar IVDs using the segmental measurements of segmental Cobb angle, A/P ratio, and IVD width. Moreover, the images obtained in standing can yield more physiologically relevant information than a slightly higher resolution scan taken in supine due to the differences in loading between supine and standing. Although some of the measurements made here could also be obtained from plain film radiographs (23, 25), the pMRI enables the possibility of assessing any potential involvement of IVD-specific pathologies (such as herniations) in the spinal structural measurements. The nature of MRI also leverages the increased signal of hydrated microstructures in the IVD, and the 3D reconstruction capabilities of the volumetric image stack could be utilized further for additional spatially-unbiased analyses.

There are several limitations to our study. First, although the field strength of this pMRI system (0.6 T) is lower than typical clinical systems (1.5 T), the higher efficiency of the field algorithm reconstructs and generates images at comparable resolutions to 1.5 T systems (13). The ICC and uncertainty analyses confirm the reliability of our measurements using the pMRI system. Our inter-observer reliability ranged from 0.68 to 0.99 and is consistent with a MRI study that reported an inter-observer ICC range of 0.73-0.95 (11). Analyses of error and uncertainty in our images reveal that the resolution can affect the measurements by 1.3%-5.4%, and thus only differences that were greater than the uncertainty range were considered to be meaningful. Second, we did not directly control for the pelvic tilt which is known to influence spinal alignment, and this may have contributed to the participants’ variations in the supine spinal alignment. Third, although we only enrolled 40 participants in this study, the power for our discriminating measurements ranged from 0.57-0.99, indicating that the sample size is adequate for most measures. Only the L3/L4 A/P ratio was powered below 0.80 at 0.57.

This study is the first to examine the differences in regional and segmental measurements between supine and standing positions in young, back-healthy participants using a pMRI system. Further, we have developed and validated a method using the pMRI system to evaluate changes in spinal measurements. Our key finding here is that the individual IVD levels within a healthy spine adapt to loaded positions non-uniformly. These findings highlight the importance of acquiring images in standing as they are different than the traditional MRI images in supine. Observing IVDs in standing and comparing symptomatic and asymptomatic individuals can be more informative of IVD function and help elucidate the discordance between imaging findings and LBP.

## Acknowledgements

The authors gratefully acknowledge Dave Magner, Travis Gould, Nicholas Woerther and the staff at The Open Upright MRI of Missouri for providing technical assistance with the imaging.

